# Classification of psychedelics and psychoactive drugs based on brain-wide imaging of cellular c-Fos expression

**DOI:** 10.1101/2024.05.23.590306

**Authors:** Farid Aboharb, Pasha A. Davoudian, Ling-Xiao Shao, Clara Liao, Gillian N. Rzepka, Cassandra Wojtasiewicz, Jonathan Indajang, Mark Dibbs, Jocelyne Rondeau, Alexander M. Sherwood, Alfred P. Kaye, Alex C. Kwan

**Author notes:** Correspondence to Alex Kwan, Ph.D., Room 111 Weill Hall, 526 Campus Road, Ithaca, NY, 14853, United States. These authors contributed equally to the work (Farid Aboharb, Pasha A. Davoudian).

## Abstract

Psilocybin, ketamine, and MDMA are psychoactive compounds that exert behavioral effects with distinguishable but also overlapping features. The growing interest in using these compounds as therapeutics necessitates preclinical assays that can accurately screen psychedelics and related analogs. We posit that a promising approach may be to measure drug action on markers of neural plasticity in native brain tissues. We therefore developed a pipeline for drug classification using light sheet fluorescence microscopy of immediate early gene expression at cellular resolution followed by machine learning. We tested male and female mice with a panel of drugs, including psilocybin, ketamine, 5-MeO-DMT, 6-fluoro-DET, MDMA, acute fluoxetine, chronic fluoxetine, and vehicle. In one-versus-rest classification, the exact drug was identified with 67% accuracy, significantly above the chance level of 12.5%. In one-versus-one classifications, psilocybin was discriminated from 5-MeO-DMT, ketamine, MDMA, or acute fluoxetine with >95% accuracy. We used Shapley additive explanation to pinpoint the brain regions driving the machine learning predictions. Our results support a novel approach for characterizing and validating psychoactive drugs with psychedelic properties.

## INTRODUCTION

Psychedelics include classic serotonergic psychedelics, such as psilocybin and 5-methoxy-*N,N*-dimethyltryptamine (5-MeO-DMT), and related psychoactive compounds, such as ketamine and 3,4-methylenedioxymethamphetamine (MDMA). These compounds have recently gained widespread interest as potential therapeutics for neuropsychiatric disorders^1, 2^. Psilocybin with psychological support is under active investigation as a treatment for major depressive disorder and treatment-resistant depression^3, 4, 5, 6, 7^. Subanesthetic ketamine has long been studied for its efficacy for treating depression^8, 9, 10^ and post-traumatic stress disorder (PTSD)^11^. The research efforts culminated in the approval of esketamine nasal spray by the FDA in the United States for treatment-resistant depression^12, 13^. Finally, MDMA-assisted psychotherapy has undergone phase III clinical trials for the treatment of moderate to severe PSTD^14, 15^. The clinical relevance has sparked intense interest in understanding the shared and distinct aspects of these compounds’ mechanisms of action.

Beyond the known psychedelics, there is also growing excitement for synthesizing novel psychedelic-inspired analogs that can be new chemical entities for therapeutics^16, 17, 18^. Ideally, the novel compounds would retain therapeutic effects while improving pharmacokinetics, minimizing perceptual effects, and eliminating cardiovascular risks. A major roadblock in this pursuit lies in developing screens that can filter thousands of psychedelic-inspired analogs to a manageable list of the most promising compounds for further in-depth characterizations. Currently, most screens operate at the molecular or behavioral level. At the molecular level, candidate compounds can be docked *in silico* with the structure of the 5-HT_2A_ receptor, followed by biochemical measurements of receptor engagement and activation of downstream G-protein and beta-arrestin pathways. This target-based approach has yielded exciting leads^19, 20, 21, 22^, but assumes that the 5-HT_2A_ receptor is the key mediator of the therapeutic effect, which has not been proven conclusively. At the behavioral level, candidate compounds may be tested in animals for defined phenotypes. Simple characterizations such as changes in animal movement patterns may be automated to increase throughput and accuracy^23, 24^. However, more complex behavioral assays relevant for depression suffer from limitations including poor construct validity and weak predictive power for drug efficacy in humans^25^.

The development of a new screening method may complement current molecular and behavioral approaches to accelerate preclinical drug discovery. Classic psychedelics and ketamine share the ability to enhance neural plasticity in the brain^26^, as evidenced by the rapid and persistent growth of dendritic spines in the rodent medial frontal cortex after a single dose of ketamine^27, 28^, psilocybin^29^, and related serotonergic receptor agonists^30, 31, 32, 33^. A promising approach may thus focus on quantifying indicators of neural plasticity in native brain tissues. To this end, immediate early genes are activated in a cell in response to increased firing activity or an external stimulus^34^. The immediate early genes are a key part of neural plasticity, because they enable neurons to adapt to stimuli by regulating gene expression, which is crucial for protein synthesis that are needed for synaptic modifications and learning^35, 36^. Taking classic psychedelics as an example, drug administration induces robust increases in the expression of immediate early genes^37, 38^, including c-Fos, that can be detected starting in as few as 30 minutes in multiple brain regions^39, 40^. More recently, technological advances in tissue clearing, light sheet fluorescence microscopy, and automated detection of nuclei have enabled high-throughput mapping of the expression of immediate early genes such as c-Fos in the whole mouse brain^41, 42^. We and others have applied this method to characterize the impact of psilocybin and ketamine^43, 44, 45^, joining a rapidly growing number of studies using brain-wide imaging of fluorescence signals to study drugs^46, 47, 48, 49, 50, 51, 52, 53, 54, 55, 56, 57^. Although these early studies have provided valuable biological insights, only one or two drugs were typically included in each study thus far. Developing the method as a drug screen requires evaluating its feasibility and accuracy on a larger panel of compounds.

In this study, we measured brain-wide c-Fos expression in male and female mice for 8 drug conditions, including a variety of psychedelics, related psychoactive compounds, and vehicle control. We developed a pipeline for analysis and classification based on explainable machine learning, determining performance in one-versus-rest and one-versus-one classification tasks. We implemented Shapley additive explanation to interpret the machine learning models to identify the brain regions driving the classifications. Collectively the results demonstrate brain-wide imaging of immediate early gene expression as a promising approach for preclinical drug discovery.

## RESULTS

### Psychedelics and related drugs in the study

For this study, we evaluated 8 drug conditions: psilocybin (PSI, 1 mg/kg, i.p., single dose), ketamine (KET, 10 mg/kg, i.p., single dose), *5*-methoxy-*N,N*-dimethyltryptamine (5-MeO-DMT or 5MEO, 20 mg/kg, i.p., single dose), 6-fluoro-*N,N*-diethyltryptamine (6-fluoro-DET or 6-F-DET, 20 mg/kg, i.p., single dose), 3,4-methylenedioxymethamphetamine (MDMA, 7.8 mg/kg, i.p., single dose), acute fluoxetine (A-SSRI, 10 mg/kg, i.p., single dose), chronic fluoxetine (C-SSRI, 10 mg/kg, i.p., one dose every day for 14 days), and saline vehicle (SAL, 10 mL/kg, i.p., single dose) (**Fig. 1a**).

**Fig. 1.**
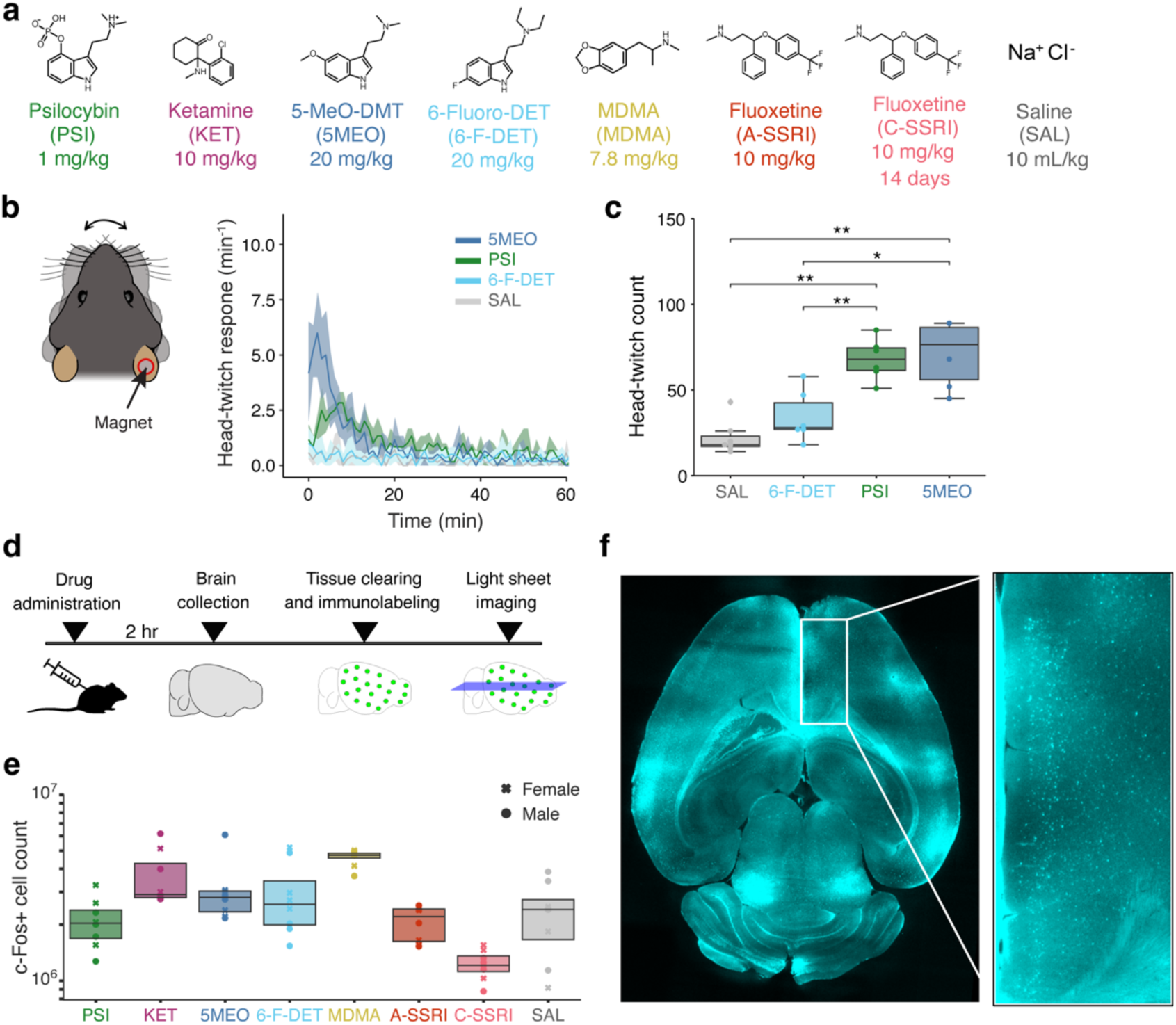
Imaging brain-wide c-Fos expression at cellular resolution following drug administration. **a.** Chemical structures for the 8 conditions included in this study: psilocybin (PSI), ketamine (KET), 5-MeO-DMT (5MEO), 6-fluoro-DMT (6-F-DET), MDMA, acute fluoxetine (A-SSRI), chronic fluoxetine (C-SSRI, daily for 14 days), and saline vehicle (SAL). **b.** Time course of head-twitch response following the administration of 5-MeO-DMT, psilocybin, 6-fluoro-DET, or saline vehicle. Line, mean. Shading, 95% confidence interval based on 1000 bootstraps. N = 3 males and 3 females for each drug, except 4 males and 3 females for saline. **c.** Box plot of the total number of head twitches detected within a 2-hour period after drug administration. Wilcoxon rank-sum test. *, *P* < 0.05, **, *P* < 0.01. **d.** Experimental timeline. **e.** Box plot of the total number of c-Fos+ cells in the brain for each drug condition. Cross, female individual. Circle, male individual. N = 64 mice, including 4 males and 4 females for each drug. **f.** An example of the fluorescence images of c-Fos+ cells in the mouse brain for a psilocybin-treated mouse acquired by light sheet fluorescence microscopy. Inset, magnified view of the dorsal anterior cingulate cortex. For **b** and **c**, the psilocybin and saline vehicle data had been shown in a prior study^33^.

We elected to investigate these compounds for several reasons. Psilocybin is a classic psychedelic that acts on the 5-HT_2A_ receptor. Psilocybin stands at the forefront of ongoing late-stage clinical trials evaluating psychedelics’ efficacy for treating depression^3, 4, 5, 6, 7^. Ketamine is primarily a NMDA receptor antagonist^58^. Despite the distinct molecular targets, ketamine and psilocybin have similarities in their plasticity-promoting action and behavioral effects^59, 60^, making ketamine an intriguing compound to contrast with psilocybin. The doses and route of administration for psilocybin and ketamine were chosen based on prior studies showing behavioral effects in mice^29, 61^.

5-MeO-DMT is a classic serotonergic psychedelic in the same tryptamine chemical class as psilocybin^16^. There is clinical interest in evaluating 5-MeO-DMT as a treatment for depression^62,63^. At a dose of 20 mg/kg in mice, 5-MeO-DMT induces head-twitch response and evokes structural rewiring in the mouse medial frontal cortex^33^. Compared to psilocybin, 5-MeO-DMT is shorter-acting and has higher affinity for the 5-HT_1A_ receptor than for the 5-HT_2A_ receptor. Thus 5-MeO-DMT serves as a useful case of another tryptamine psychedelic with distinct pharmacokinetics and receptor target profile. 6-fluoro-DET is also a tryptamine like psilocybin and 5-MeO-DMT. Although bioavailable in the brain and a 5-HT_2A_ receptor agonist^64, 65^, 6-fluoro-DET induces autonomic effects without causing perceptual changes in humans^66^. Therefore, it has been used as an active, non-hallucinogenic control in a clinical study^67^. Concordantly, 6-fluoro-DET provided ineffective as a substitute compound for rats trained to discriminate LSD or 2,5-dimethoxy-4-iodoamphetamine (known as DOI)^64, 68^. To corroborate these prior results, we measured the effect of 6-fluoro-DET on head-twitch response in mice using magnetic ear tags for automated detection of head movements. Our results showed that, unlike 1 mg/kg psilocybin and 20 mg/kg 5-MeO-DMT which elicited robust head-twitch responses^33^, mice administered with 20 mg/kg 6-fluoro-DET were not statistically different from controls (**Fig. 1b, c**). Our study adds to other recent studies^20, 21^ that included 6-fluoro-DET as a non-hallucinogenic tryptamine for comparison. The dose of 6-fluoro-DET was chosen to match the dose of 5-MeO-DMT.

MDMA is different from psilocybin: it is a member of the phenethylamine chemical class and has distinct pro-social and euphoric qualities^69^. MDMA can act on monoamine transporters to enhance release and inhibit reuptake of neuromodulators including serotonin, thus it has been characterized as an entactogen rather than a classic psychedelic^70^. MDMA holds clinical relevance particularly for PTSD^14, 15^. We selected a dose of 7.8 mg/kg for MDMA based on prior work showing that this dose facilitates fear extinction learning in mice^71^. Fluoxetine is a commonly prescribed antidepressant that is a selective serotonin reuptake inhibitor (SSRI). Clinical interest lies in understanding the relative efficacies of SSRIs versus psilocybin^4^ and whether ketamine or psilocybin is suitable for treatment-resistant depression^5, 12, 13^. SSRIs require chronic administration to exert therapeutic effects, therefore likely engage a mechanism of action distinct than that of psilocybin and ketamine. For these reasons, we included acute and chronic fluoxetine for this study. We chose a dose of 10 mg/kg, which was used for acute and chronic administration of fluoxetine in mice previously^72, 73^. Control animals received a single injection of saline vehicle.

### Light sheet fluorescence imaging of cellular c-Fos expression

For each of the 8 drugs, we tested 4 male and 4 female C57BL/6J mice, totaling 64 animals for the entire data set. Brains were collected 2 hours after the administration of the single dose or 2 hours after the administration of the last dose for the chronic fluoxetine condition (**Fig. 1d**). The 2-hour interval was chosen assuming drug penetrance to the brain by 0.5 hours and peak c-Fos expression after an additional 1.5 hours^74^. Brains were processed for tissue clearing and c-Fos immunohistochemistry (see **Methods**). Light sheet fluorescence microscopy was used to image each brain at a resolution of 1.8 µm per pixel in the x- and y-axis and at 4 µm intervals in the z-axis, which allowed for sampling of all cells in the entire brain without any gap. The images were analyzed using neural nets for automated detection of fluorescent puncta corresponding to c-Fos+ cells (see **Methods**). The number of c-Fos+ cells detected in each brain for each condition is presented in **Figure 1e**. An example image collected from a mouse administered with psilocybin is shown in **Figure 1f**.

To investigate the regional distribution of c-Fos+ cells, we aligned the images of each brain to the Allen Brain Atlas and segmented the images into summary structures based on the Allen Mouse Brain Common Coordinate Framework^75^ (see **Methods**; **Supplementary Table 1**). The number of c-Fos+ cells in each brain region for all animals is provided in **Supplementary Table 2**. To visualize the entire data set, we normalized the c-Fos+ cell count in each brain region by the total number of c-Fos+ cells of each brain and by the spatial volume of the brain region. **Figure 2** is a heatmap of the resulting c-Fos+ cell density for all the samples. We observed that c-Fos+ cell density was generally high in the isocortex, olfactory area, hippocampal area, striatum and pallidum, and thalamus, whereas expression was lower in the midbrain and hindbrain, and cerebellum. There were individual differences across samples from the same drug, but also notable contrasts across different drugs. This begets questions such as: How does the individual variability compare with the differences across drugs? How well can whole-brain c-Fos maps be used to discriminate the different drugs?

**Fig. 2.**
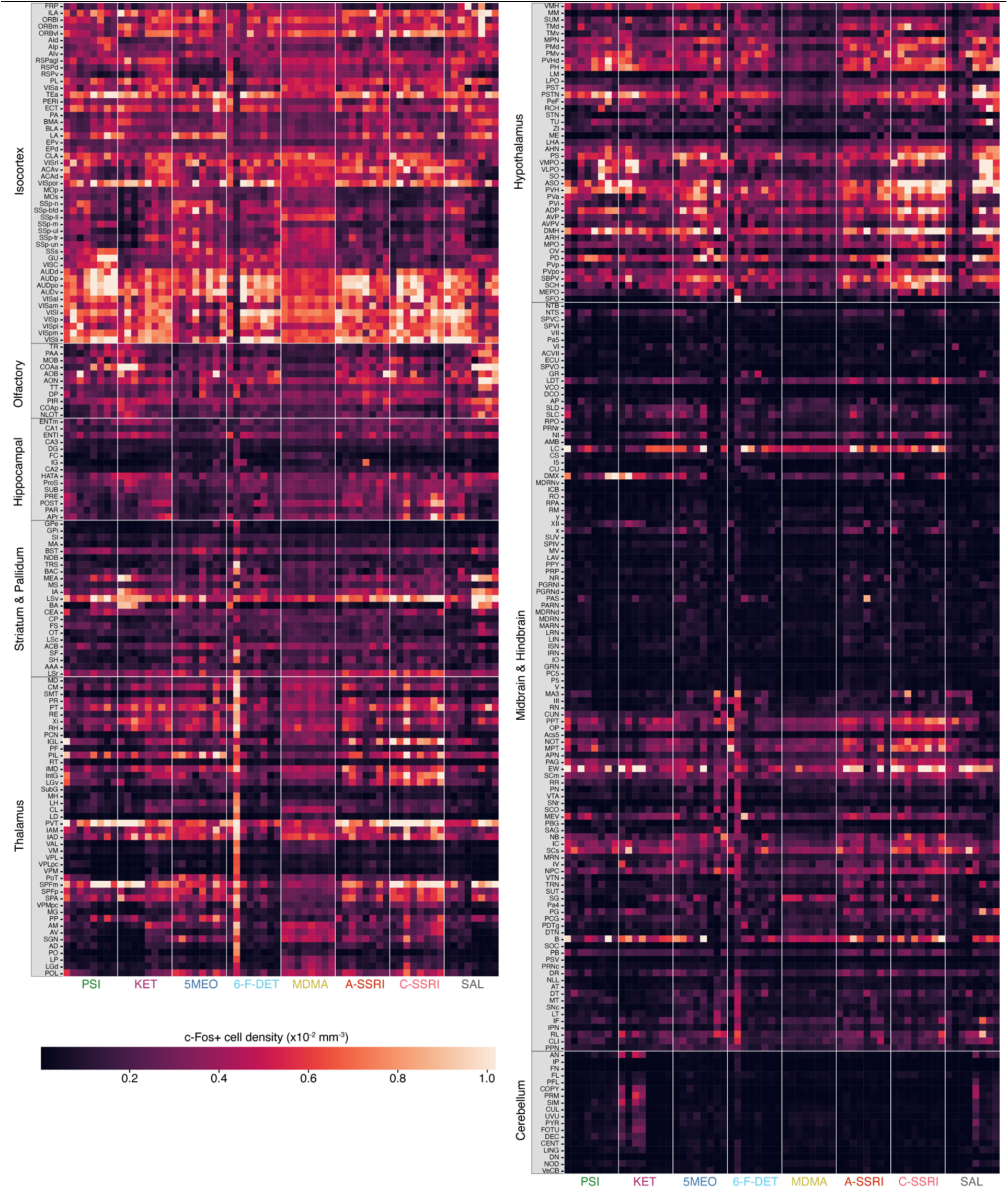
c-Fos+ cell density listed by brain region for all samples by drugs. The c-Fos+ cell density was defined as the c-Fos+ cell count in each brain region divided by the total number of c-Fos+ cells in each brain and the spatial volume of the brain region. The pixels in the heatmap are positioned by brain region (row) and animal grouped by drug (column). The intensity of the pixel is pseudo-colored by the value of the c-Fos+ cell density. The brain regions including acronyms and other details are provided in **Supplementary Table 1**.

### Machine learning pipeline for classifying drugs based on brain-wide c-Fos distribution

To answer these questions, we developed a pipeline for quantitative comparison of the brain-wide c-Fos expression data between different drug conditions. We posited that different compounds may elicit distinct regional distribution of cellular c-Fos expression that can serve as fingerprints for classifying drugs. The pipeline starts with a matrix of c-Fos+ cell counts for different brain regions from different samples (first panel, **Fig. 3a**). This matrix of c-Fos+ cell counts undergoes preprocessing, starting with normalization (dividing the c-Fos+ cell count in each region by the total c-Fos+ cell count of the brain) (second panel, **Fig. 3a**). Normalization is important because there may be batch effects across samples. The data were then processed to scale the input data to a standard range such that the values across brain regions are more comparable and amenable to fitting machine learning models (second panel, **Fig. 3a**), using Yeo-Johnson transformation (monotonic transformation of data using a power function) and robust scaling (median subtraction and interquartile range scaling). We will herein refer to the values after this preprocessing step as the c-Fos scores.

**Fig. 3.**
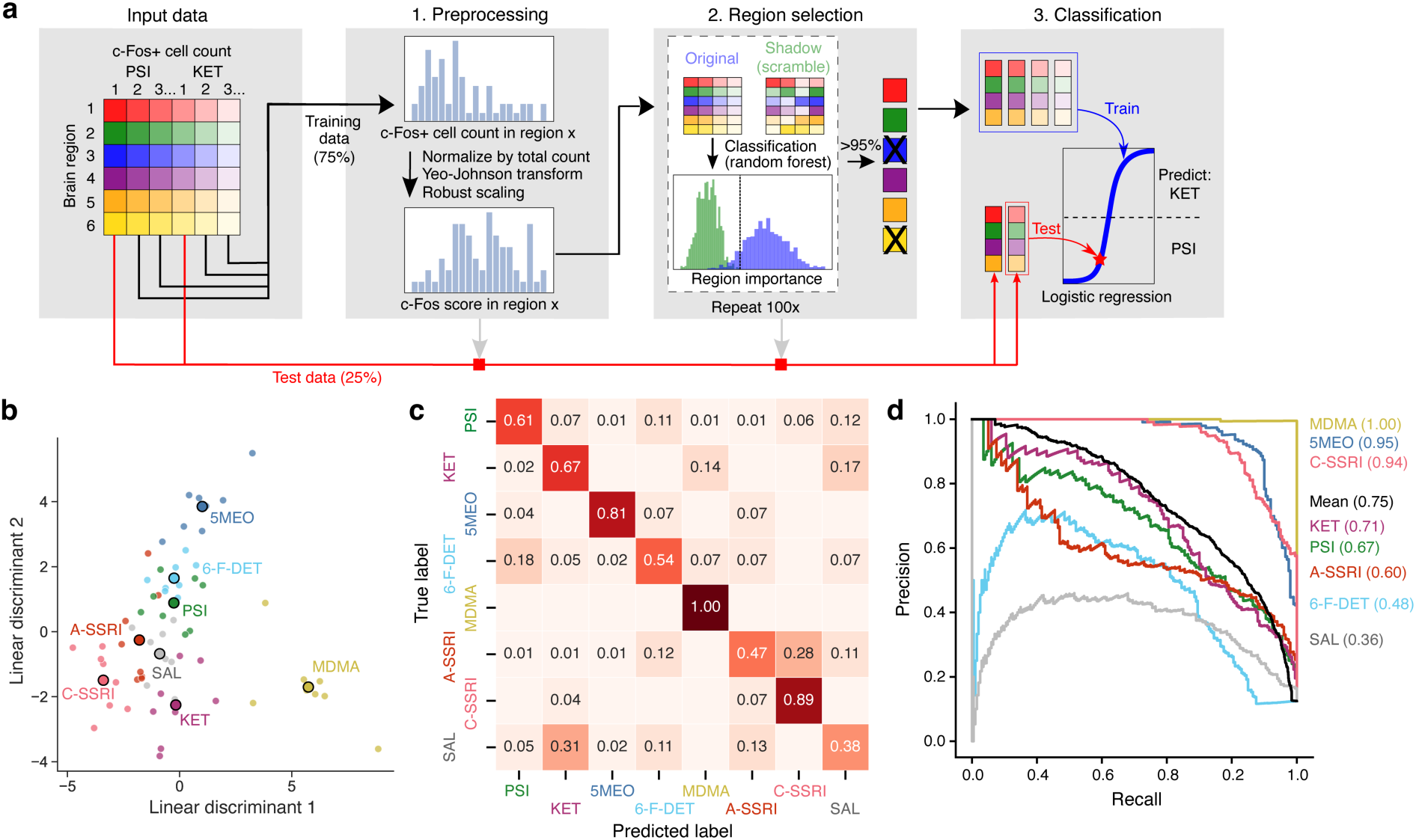
A machine learning pipeline for drug prediction and performance of one-versus-rest classification. **a.** The pipeline consisted of three steps. First, c-Fos+ cell counts for each brain region undergo normalization, Yeo-Johnson transformation, and robust scaling, into c-Fos scores. Second, the Boruta procedure is used to select the set of informative brain regions. Third, c-Fos scores from this set of brain regions were used to fit a ridge logistic regression model. For each iteration, 75% of the data in each drug condition were used for region selection and training through the three steps, and the remaining 25% of the data were withheld initially, but then processed and tested with the ridge logistic regression model. The entire process was iterated using different splits of the data for 100 times. **b.** Linear discriminant analysis of the c-Fos scores to visualize the data in a low dimensional space. **c.** The confusion matrix showing the mean proportion of predicted labels for each of the true labels across all splits. **d.** The composite precision-recall curves for each drug condition across all splits and the grand average across all drugs. The values in parentheses are the area under the precision-recall curve for the compounds.

Next, we adapted the Boruta feature selection procedure^19^ to determine which brain regions to include for model fitting and testing (third panel, **Fig. 3a**). The Boruta procedure is a permutation-based method for determining feature importance. It starts by creating “shadow features”: for example, if the data contains 48 c-Fos scores for brain region 1 for various conditions, then the corresponding shadow feature will be those same 48 c-Fos scores with scrambled drug labels. Shadow variants were created for all brain regions to create the expanded Boruta dataset. A random forest classifier was built using this Boruta dataset to determine a feature-importance value for each brain region. If a brain region has a higher feature-importance value than the largest feature-importance value from shadow features, then brain region 1 is a “hit”. This permutation process is iterated 100 times. Given that each brain region can achieve only one of two outcomes (hit or no hit) in each iteration, the distribution of outcomes across all iterations is a binomial distribution, and a brain region is included by the statistical criterion of exceeding 95^th^ percentile of the binomial distribution. Why Boruta? We used the Boruta procedure in lieu of including all brain regions, because many regions likely contribute little or nothing towards differential drug action and their inclusion in the model would increase noise and lead to overfitting. A distinctive advantage of Boruta is that brain regions do not compete with each other, but rather with the shadows. As a result, the number of brain regions selected by Boruta is not pre-determined but instead dictated by the data as needed.

For the last step, the c-Fos scores from the selected brain regions are used to construct a ridge logistic regression model (fourth panel, **Fig. 3a**). The entire pipeline is evaluated using 4-fold splits, where 75% of the data in each drug condition was used to train and fit the model, while the remaining 25% of the data is used to test the model. Importantly, we emphasize that we used only the training data to optimize the preprocessing parameters, run feature selection, and construct regression model. The same optimized preprocessing parameters and selected features were then later applied for the test data, ensuring no data leakage. The splits were repeated 100 times to evaluate the prediction accuracy of the pipeline.

### One-versus-rest classification shows drug prediction accuracy well above chance

We performed a linear discriminant analysis on the c-Fos scores of all 64 samples, just after the preprocessing step. We plotted the data for the top two linear discriminants (**Fig. 3b**). This visualization clearly shows that the differences in c-Fos scores across drugs are more separable than the differences in c-Fos scores across samples within the same drug condition. Drugs that alter the serotonergic tone via different mechanisms of action are positioned differently along the first linear discriminant. By contrast, 5-MeO-DMT, 6-fluoro-DET, and psilocybin are separable along the second linear discriminant.

We first tested the pipeline with the entire data set and asked the models to predict the exact drug condition. The confusion matrix shows how the predicted drug labels compared with the true drug labels (**Fig. 3c**). Because there were 8 conditions, the chance-level accuracy was 12.5% (1 out of 8). We found that the model was the most accurate at identifying the MDMA, chronic fluoxetine, and 5-MeO-DMT samples, with 100%, 89%, and 81% accuracy respectively. Performance for other conditions were lower, yielding an overall mean accuracy of 67% for all drugs. Performance was the lowest for saline and acute fluoxetine at 38% and 47% respectively. Our interpretation for the low-performance conditions is that tradeoffs must be made to solve this 8-way classification problem. The machine learning model uses the cross-entropy loss function, which seeks to maximize the probability of labeling training data correctly across the entire training set, rather than drawing boundaries in a one-vs-rest fashion. In this global approach, individual decision boundaries may be placed in a way which under performs on one label, such as saline, while leading to a greater improvement on others. In other words, the model was fitted with the goal of maximizing the overall mean classification accuracy, which was not necessarily the most ideal for distinguishing any one specific condition such as saline. Nevertheless, the mean accuracy of 67% was still substantially higher than chance level of 12.5%.

Confusion matrices are calculated based on a single decision threshold, which may exaggerate true positive rate for one drug type at the expense of more false positives for another drug type. To understand our model performance from a different perspective, we plotted precision-recall curves (**Fig. 3d**). These curves consider performance across all possible decision thresholds and summarize the results in terms of precision (true positives relative to false positives) and recall (true positives relative to false negative). The perfect classifier would have an area under the precision-recall curve (precision-recall AUC) of 1. Across all drugs, the pipeline yielded a mean precision-recall AUC value of 0.75. This is well above the theoretical chance-level of 0.125 for 1 out of 8 drugs and the empirical chance-level of 0.12 calculated with shuffled data. The performance based on precision-recall AUC for predicting different drugs corresponds in rank order to the accuracy in the confusion matrix. Overall, these results provide evidence that brain-wide c-Fos maps can be leveraged to identify the exact drug administered out of a panel of related psychoactive compounds.

A likely use case for the pipeline is to determine how a novel chemical entity may be positioned in the pharmacological space based on the c-Fos expression pattern. To simulate this scenario, we performed a leave-one-drug-out analysis, in which we trained a model using 7 conditions (psilocybin, ketamine, 5-MeO-DMT, MDMA, acute fluoxetine, chronic fluoxetine, and saline), but then tested it on all conditions including 6-fluoro-DET. We found that 6-fluoro-DET was most frequently classified as psilocybin at 44% chance but could also be detected as saline at 29% chance (**Fig. S1**), which is in general agreement with 6-fluoro-DET being a non-hallucinogenic 5-HT_2A_ receptor agonist.

### One-versus-one classification suggests a small list of brain regions drives drug prediction

We reasoned that one-versus-one classification, where the machine learning pipeline solves a binary problem of deciding between two drugs (**Fig. 4a**), may provide deeper insights into the factors that distinguish specific drug classes. Given the prominence of psilocybin in clinical trials and drug discovery, we were particularly interested in comparisons between psilocybin and other conditions that differ in serotonergic receptor affinities (5-MeO-DMT), mechanism of action (MDMA, acute fluoxetine, ketamine), or hallucinogenic potency (6-fluoro-DET). We trained the same machine learning pipeline using subsets of data involving only two or three drugs. The binary classifiers achieved near-perfect accuracy reflected by precision-recall AUC values at or exceeding 0.90, with the notable exception of psilocybin versus 6-fluoro-DET which had a precision-recall AUC of 0.59 (**Fig. 4b**). The difficulty in discerning between a classic serotonergic psychedelic and the non-hallucinogenic 5-HT_2A_ receptor agonist extended beyond psilocybin: 5-MeO-DMT versus 6-fluoro-DET as well as psilocybin and 5-MeO-DMT versus 6-fluoro-DET also yielded modest precision-recall AUC values at 0.80 and 0.57 respectively, relative to chance level of 0.5 for one-versus-one classifications. These results suggest that brain-wide cellular c-Fos expression is effective at discriminating between exemplars from different drug classes, such as a classic psychedelic versus an entactogen, a classic psychedelic versus a dissociative, and a classic psychedelic versus SSRI. It also effectively distinguishes between the two classic psychedelics psilocybin and 5-MeO-DMT. However, the prediction is less reliable for the specific problem of predicting a non-hallucinogenic 5-HT_2A_ receptor agonist relative to a classic psychedelic.

**Fig. 4.**
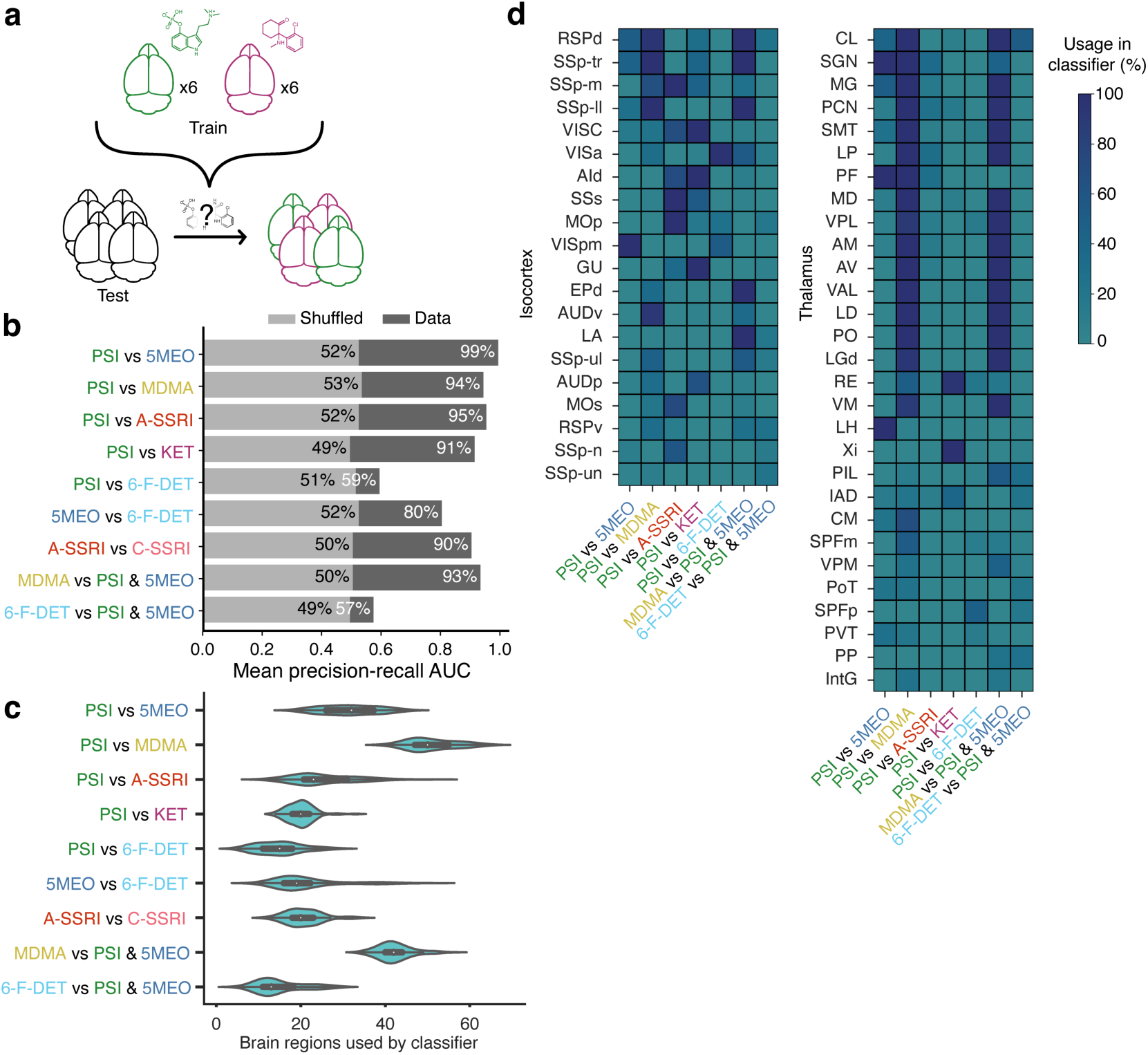
Performance of one-versus-one classification. **a.** Schematic illustrating the one-versus-one classification problem. **b.** The mean area under the precision-recall curve across all splits for different binary classifiers. Dark gray, real data. Light gray, shuffled data. **c.** The number of brain regions selected via the Boruta procedure for inclusion in the regression model. **d.** Heatmaps showing the fraction of splits when a cortical (left) or thalamic (right) region was included in the regression model. The regions are sorted based on usage in all classifiers. Regions that were included in <75% of the splits across all conditions are not shown.

As mentioned, a feature of the Boruta procedure is that a different number of regions may be included depending on the data and the desired classification. Indeed, there were differences in the brain regions chosen for the various drug prediction problems and different training and testing splits of the same data (**Fig. 4c**). Most classifiers relied on <35 brain regions for drug prediction, except for the two comparisons involving MDMA which included around 40 - 70 brain regions. Furthermore, we plotted how often various cortical and thalamic regions were selected by the machine learning models (**Fig. 4d**). Regions such as retrosplenial areas (RSPd, RSPv), somatosensory areas (SSp-m, SSp-tr, SSp-II), and lateral networks (VISC, AId) were included often, but different classifiers relied on them to different extents. We will explore the importance of specific brain regions quantitatively in the next section using Shapley additive explanation. Many thalamic regions were consistently included in comparisons involving MDMA, which contributed to the higher total number of brain regions used by classifiers when MDMA was involved. Overall, the results suggest that one-versus-one drug classifications based on brain-wide c-Fos expression is highly accurate, with the machine learning models only needing data from a small number of brain regions to produce the prediction.

### Using Shapley additive explanation to highlight key brain regions driving drug prediction

A brain region selected by Boruta in the pipeline suggests that it is informative, yet it does not communicate the importance of its contribution to the final prediction. To better understand how the c-Fos scores in individual brain regions contribute to decisions in one-versus-one drug classifications we used Shapley additive explanation (SHAP) (**Fig. 5a**). SHAP uses a game-theoretical approach to determine how the brain regions contribute to driving the machine learning regression model from a starting base value to the final output value for decision^21^. To illustrate, we present the force plot of two test brain samples in one of our cross-validation splits (**Fig. 5b**). The top half of the plot shows the c-Fos scores in selected brain regions for the sample of psilocybin and their additive contributions to the decision. In this instance, regions such as posteromedial visual area (VISpm, c-Fos score = 0.44) and lateral habenula (LH, c-Fos score = -0.78) were among the drivers leading to an overall positive SHAP value to predict psilocybin. The posteromedial visual area is located between the primary visual cortex and retrosplenial cortex^76^ and has been suggested to mediate visual information between the neighboring regions^77^. Lateral habenula neurons had spiking activity associated with undesirable outcomes^78, 79^, which is consistent with their posited role in mediating depression-related symptoms^80^ and contributing to antidepressant response^81^. Intriguingly, another driver was the parafascicular nucleus (PF, c-Fos score = -1.74), which is implicated in arousal and head movements^82^. By contrast, the c-Fos scores in the same set of selected brain regions sums to an overall negative SHAP value for the 5-MeO-DMT sample, providing the basis for the correct prediction in this case. Across all splits tested for the psilocybin-versus-5-MeO-DMT comparison, we identified regions that were included in >75% of the machine learning models, and then ranked these regions by mean SHAP value difference, which highlight the brain regions most responsible for driving the classification (**Fig. 5c, d**).

**Fig. 5.**
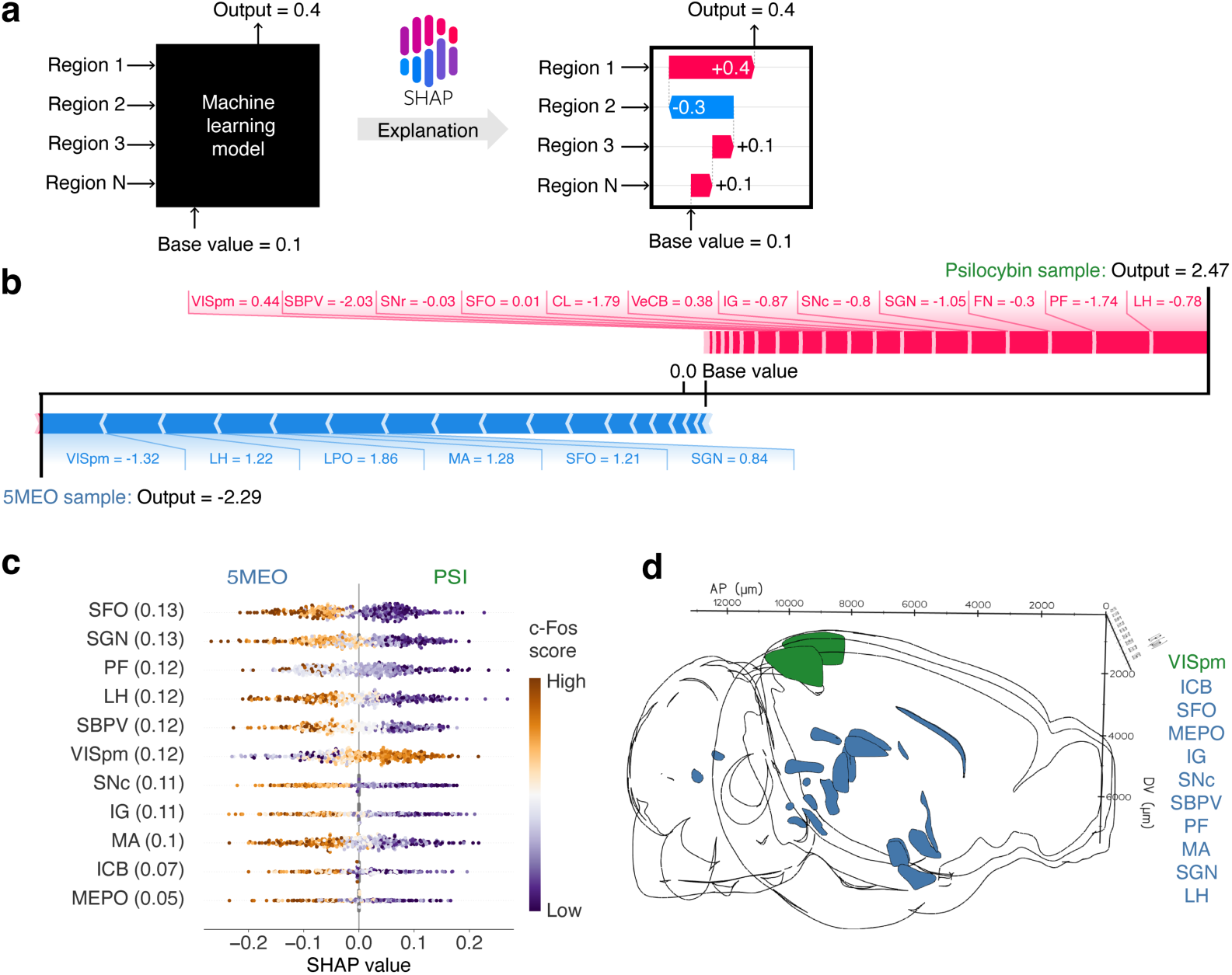
Shapley additive explanation for identifying brain regions driving the prediction of 5-MeO-DMT from psilocybin. **a**. Diagram illustrating the concept behind SHAP values. The ridge regression model is akin to a black box that takes c-Fos scores as inputs to produce a prediction. SHAP values can be computed to quantitatively assess how strong and in what direction the c-Fos score of each brain region contributes to the prediction. **b**. Example force plots for a psilocybin sample and a 5-MeO-DMT sample from one split, illustrating how actual c-Fos scores of brain regions add to shift the model’s output from the base value to the final value. **c**. Plot relating a region’s c-Fos scores to the SHAP values across individual splits of the 100 iterations for the 5-MeO-DMT-versus-psilocybin classification. Brain regions were shown only if they were used by >=75% of the splits and listed in rank order by the absolute value of the mean difference in SHAP values between the two drug conditions. The values in parentheses are the absolute value of the mean difference in SHAP values between the two drug conditions. **d**. Visualization of the brain regions included in **c**, color coded according to the compound which evoked higher c-Fos score in the region.

We also analyzed other one-versus-one classification problems using Shapley additive explanation. For MDMA versus psilocybin, there was a longer list including 32 brains regions that were used in at least 75% of the cross-validation splits (**Fig. 6a, b**). Half of these regions (16/32) were in the thalamus. Given the larger number of regions in each model, the SHAP value differences tended to be smaller, because there is redundancy in the information provided by the regions.

**Fig. 6.**
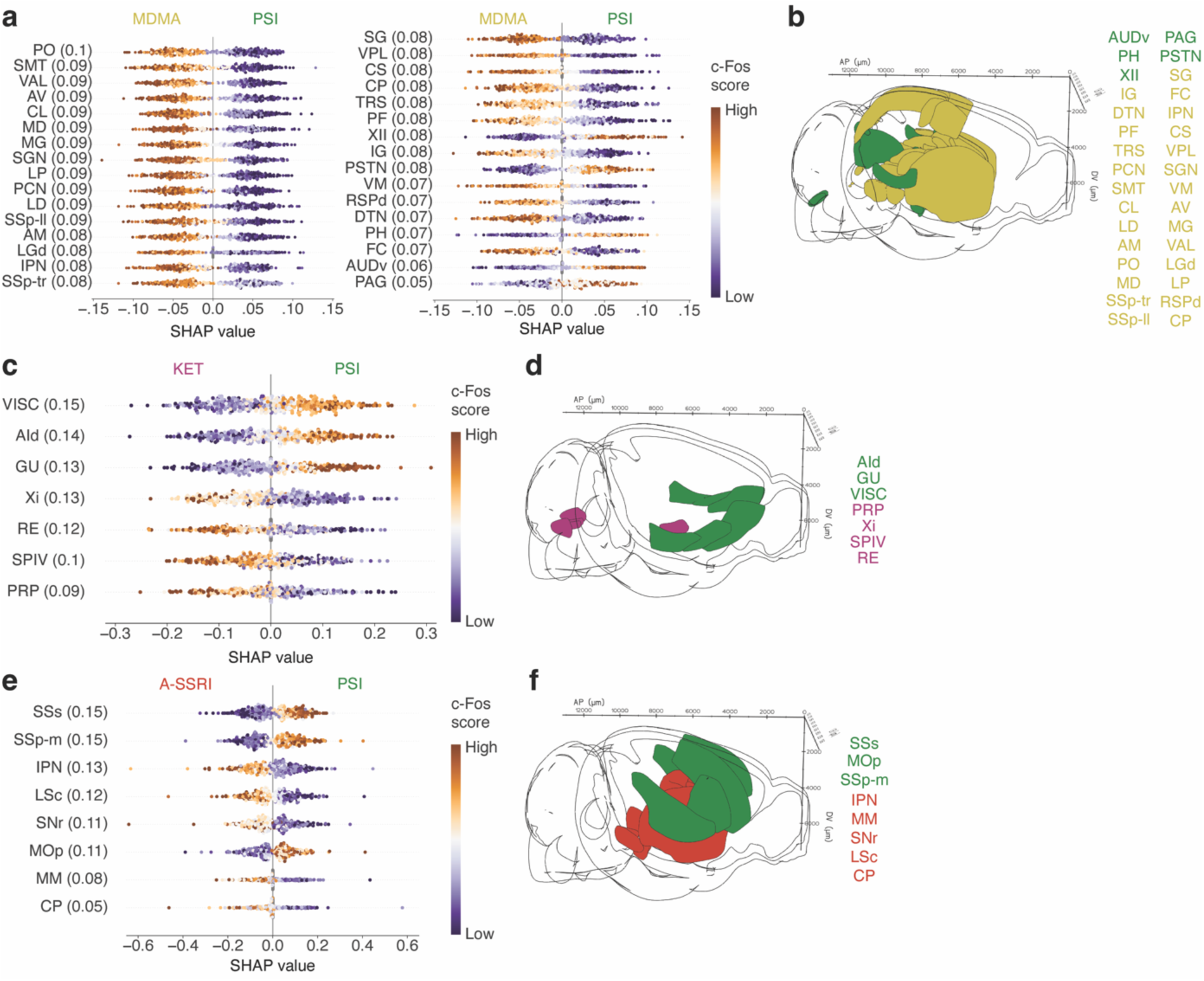
Brain regions driving the prediction of MDMA, ketamine, or fluoxetine from psilocybin. **a, b.** Similar to Fig. 5c, d for MDMA-versus psilocybin classification. **c, d.** Similar to Fig. 5c, d for ketamine-versus psilocybin classification. **e, f.** Similar to Fig. 5c, d for acute fluoxetine-versus psilocybin classification.

For ketamine versus psilocybin, the top 5 regions that were consistently included in >96% of the cross-validation splits and had the highest SHAP value differences were the visceral area (VISC), gustatory area (GU), dorsal agranular insular area (AId), xiphoid thalamic nucleus (Xi), and nucleus of reuniens (RE) (**Fig. 6c, d**). VISC and GU have direct connections to AId, all of which are part of the lateral subnetworks of the mouse neocortex^83, 84^. The mouse insular cortex contains various cell types that express an abundance of 5-HT_2A_ and 5-HT_1A_ receptors^85^, which may predispose it to stronger activation by psilocybin. Indeed, the higher c-Fos scores in these lateral cortical regions informed the model to predict psilocybin. Of note, the insular cortex is considered a core region in the mouse homolog of the salience network^86, 87^, which has been implicated in mood regulation and depression in humans^88^. Xi and RE are part of the midline thalamus, which receives visual inputs to mediate behavioral responses to threat^89^.

Interestingly, higher c-Fos scores in these midline thalamic regions are routinely used by the machine learning models to predict ketamine.

Finally, we also plotted SHAP value differences comparing acute fluoxetine and psilocybin (**Fig. 6e, f**). Here, the strongest differences were detected by in regions involved in somatosensation and motor control, including cortical somatosensory regions (SSs, SSp-m), primary motor cortex (MOp), substantia nigra (SNr), and caudoputamen (CP). These effects may relate to the previously noted effects of psychedelic on the integration of tactile sensory inputs^90^. Other implicated regions are the interpeduncular nucleus (IPN) and medial mammillary nucleus (MM), which are deep midbrain regions that are component of the limbic midbrain circuitry with long-range connections to habenula, amygdala, and hippocampus.

## DISCUSSION

In this study, we evaluated the possibility of using whole-brain imaging of cellular c-Fos expression for drug classification. We developed a machine learning pipeline with key features including adapting the statistical Boruta procedure to select informative brain regions and using Shapley additive explanation to identify features that drive the classifications. We tested the approach using 64 mice that were administered with a panel of psychedelics and related psychoactive drugs. The results demonstrated high accuracy in various one-versus-rest and one-versus-one classification problems, supporting the utility of the approach for preclinical drug discovery. For dissemination, the data and code are available at a public repository.

Immunohistochemistry can be influenced by factors such as fixation method, incubation time, antibody quality, and antigen retrieval techniques. Consequently, the c-Fos antibody staining can differ from sample to sample. Here, the issue of inter-sample variability was mitigated by not using the absolute c-Fos+ cell counts for analysis, but instead using the proportional distribution in each brain region by dividing c-Fos+ cell counts in each region by the total count in each brain. For instance, if the entire brain was stained poorly and the total c-Fos+ cell count is low, the proportion distribution should remain unchanged. This normalization step is possible when whole-brain data is acquired via light sheet fluorescence microscopy. Experimentally, the variation in antibody staining is also reduced because active electrotransport methods were used for immunolabeling. Although the normalization step is expected to help with inter-sample variability, we note that the 64 samples were processed for imaging over 3 batches (details are provided in **Methods**), and some differences may arise from batch effects.

On average, only a small number of brain regions (∼25 brain regions, except for the two comparisons involving MDMA which included ∼50 brain regions) out of the >300 summary structures in the brain were included in the machine learning models. From our prior study comparing psilocybin and ketamine^43^, we know that both compounds induce increases in c-Fos+ expression in numerous brain regions including dorsal and ventral anterior cingulate cortex (ACAd, ACAv), prelimbic area (PL), primary visual cortex (VISp), retrosplenial cortex (RSP), mediodorsal thalamus (MD), locus coeruleus (LC), lateral habenula (LH), claustrum (CLA), basolateral amygdala (BLA), and central amygdala (CEA). These brain regions are likely important for drug action, but shared targets of ketamine and psilocybin are not helpful for distinguishing the compounds. By design, the machine learning pipeline emphasizes brain regions with c-Fos expression changes that can discriminate between drug conditions, for which we found a short list of brain regions.

We anticipate the pipeline to be useful for classifying new chemical entities. For instance, when a novel psychedelic-inspired compound is synthesized, we may predict its action in the brain by its position in the linear discriminant axes (**Fig. 3b**) and the proximity to existing drug labels (**Fig. 3c**). We simulated how such a scenario could work by fitting the pipeline with 7 compounds and testing 6-fluoro-DET as if the classifier has never seen it previously (**Fig. S1**). For the full panel of drugs tested, we show that the exact drug could be identified with mean accuracy of 67%, significantly above the chance level of 12.5%. It is instructive to ask how the pipeline’s performance compared with other approaches to classify drugs. For humans, psilocybin, ketamine, and MDMA exert comparable acute behavioral effects in metrics such as experience of unity, oceanic boundlessness, and changed meaning of percepts^69^. However, MDMA preferentially induce blissful state, whereas ketamine evokes disembodiment and psilocybin induces elementary imagery and audio-visual synesthesia^69, 91^. In one study, human participants were asked to guess the administered drug, choosing between mescaline (500 mg and 300 mg), LSD, and psilocybin^92^. The accuracy for identifying the correct drug ranged from 48% to 58% during the session and 69% to 81% after the study. For animals, there has been recent progress in capturing videos of freely moving mice and analyzing their motion using unsupervised machine learning methods. One study used motion sequencing method to investigate a larger panel of 30 psychoactive compounds and doses from a wide range of drug classes including benzodiazepines, antidepressants, antipsychotics, and stimulants (but not psychedelics and the compounds tested in the current study) to show a F1 precision-recall score of 0.62^23^. Our pipeline based on brain-wide cellular c-Fos expression and machine learning therefore performed at a level comparable to earlier methods based on human and animal behaviors.

As with any analysis pipeline, there are methodological choices that can affect the outcome, which can plague the interpretation as demonstrated in the field of neuroimaging^93^. Our codebase is available online for anyone to freely use, adapt, and test. We used a statistical method with the Boruta algorithm, rather than a strict threshold, for region selection. We were careful about data leakage, using only the training data for parameter optimization and feature selection, such that the prediction accuracy for test data would not be inflated. We implemented Shapley additive explanation to decipher the factors driving the decisions, which is a general approach that should find great utility in neuroscience^94^, and has already seen applications in behavioral classification^95^ and spike waveform analyses^96^. There are areas of improvement for the pipeline. While we opted for the simplicity of treating each brain region on its own, regions may have correlated responses to drug administration. There may be biological reasons, such as anatomical proximity or synaptic connectivity, for clustering brain regions prior to region selection, which may outperform our procedure. Network analyses may be used to explore potential correlated responses to drugs. Furthermore, the pipeline will benefit from testing a larger range of compounds including enantiomers, other drug classes, and different doses. The drugs may be administered in conjunction with a receptor antagonist and a stress or behavioral manipulation, which will all lead to a richer and more refined picture of the ‘drug space’. Finally, c-Fos is one immediate early gene. It is well characterized as an activity-dependent gene and has the advantage of nuclear labeling that permits automated detection. However, there are other immediate early genes and plasticity-related biomarkers that can provide complementary information.

Here we only demonstrated moderate throughput by performing the whole-brain imaging approach for a sample size of 64 brains. This falls short of other current screening methods, which typically involve hundreds of conditions including more compounds, different doses, and additions of antagonists for competitive assays. For whole-brain imaging, the main issue was cost, which precluded us from testing at a larger scale. At the moment, the drug injection and tissue extraction steps are straightforward. The cell counting procedure is mostly automated. However, the cost per brain is high due to tissue processing and imaging, which may drop in the future because of the rapid advances in brain clearing methods^97^ and the development of inexpensive light sheet fluorescence microscopes^98, 99^. Thus, there is hope that whole-brain imaging can become a practical method for screening drugs within the next several years.

In summary, there is intense interest in using psychedelics for the treatment of neuropsychiatric disorders. Progress hinges on knowing more about existing psychedelics and finding new psychedelic-inspired drugs with improved characteristics. However, there is currently a paucity of reliable methods to screen psychedelics and related analogs. Here we developed and characterized an approach based on whole-brain imaging of cellular c-Fos expression. We demonstrated high prediction accuracy for drug classifications using a machine learning pipeline. We expect this and other neuroscience-based approaches to play an important role for accelerating the preclinical development of psychiatric drugs.

## Acknowledgments

Psilocybin, 5-MeO-DMT, and 6-fluoro-DET were provided by Usona Institute’s Investigational Drug & Material Supply Program; the Usona Institute IDMSP is supported by Alexander Sherwood, Robert Kargbo, and Kristi Kaylo in Madison, WI. This work was supported by NIH grants R01MH121848 (A.C.K.), R01MH128217 (A.C.K.), R01MH137047 (A.C.K.); One Mind – COMPASS Rising Star Award (A.C.K.); Cornell Engineering M.D.-M.Eng. program (F.A.); NIH training grants T32GM007205 (P.A.D.), T32NS041228 (C.L.); NIH fellowship F30DA059437 (P.A.D.); Source Research Foundation student grant (P.A.D.); VA National Center for PTSD (A.P.K.); Department of Defense HT9425-23-1-0458 (A.P.K.); K08MH122733 (A.P.K.); this work was funded in part by the State of Connecticut, Department of Mental Health and Addiction Services, but this publication does not express the views of the Department of Mental Health and Addiction Services or the State of Connecticut. The views and opinions expressed are those of the authors.

## Contributions

F.A., P.A.D., and A.C.K planned the study. P.A.D., L.X.S., and C.L. performed experiments. G.N.R. and C.W. assisted with tissue processing and imaging. P.A.D. and M.D. measured head-twitch responses. F.A. and P.A.D. analyzed the data, with input from A.C.K. on the pipeline. J.I. assisted with data analysis. J.R., A.M.S., and A.P.K. contributed reagents. F.A. and A.C.K. drafted the manuscript. All authors reviewed the manuscript before submission.

## Data availability

Data and code associated with the study will be available on https://github.com/Kwan-Lab.

## METHODS

### Animals

We used adult, 8-week-old male and female C57BL/6J mice (#00064, The Jackson Laboratory). Tissues were collected and imaged in three batches. The first batch performed in August 2021 included 2 males and 2 females for psilocybin (1 mg/kg, i.p.), 2 males and 2 females for ketamine (10 mg/kg, i.p.), and 2 males and 2 females for saline (10 mL/kg, i.p.). Data from these mice were included in a previous study^12^. The second batch performed in May 2022 included 2 males and 2 females for psilocybin (1 mg/kg, i.p.), 2 males and 2 females for saline (10 mL/kg, i.p.), 4 males and 4 females for 5-MeO-DMT (20 mg/kg, i.p.), 4 males and 4 females for 6-fluoro-DET (20 mg/kg, i.p.), 4 males and 4 females for acute fluoxetine (10 mg/kg, i.p.), 4 males and 4 females for chronic fluoxetine (10 mg/kg, i.p.; daily for 14 days). The third batch performed in December 2022 included 4 males and 4 females for MDMA (7.8 mg/kg, i.p.) and 2 males and 2 females for ketamine (10 mg/kg, i.p.). All animals were housed and handled according to protocols approved by the Institutional Animal Care and Use Committee (IACUC) at Yale University and Cornell University. Tissue collection for all batches was done at Yale University, except for ketamine in the third batch that was done at Cornell University. For all batches, the brain samples were shipped for clearing and imaging at LifeCanvas Technologies (Cambridge, MA).

### Drugs

Psilocybin, 5-MeO-DMT succinate, and 6-fluoro-DET solids were obtained from Usona Institute’s Investigational Drug & Material Supply Program. We used the succinate salt form of 5-MeO-DMT^100^ (at equivalent amount to freebase 5-MeO-DMT) because it can be dissolved in saline. Ketamine hydrochloride injection vial (055853, Henry Schein; or Dechra), fluoxetine hydrochloride solid (F132, Millipore-Sigma), 3,4-MDMA hydrochloride (13971, Cayman Chemical), and saline (NDC: 0409-4888-03, Hospira) were purchased from supply vendors. Psilocybin, 5-MeO-DMT succinate, 6-fluoro-DET, MDMA, and fluoxetine were prepared by dissolving powders into saline. Ketamine was prepared by diluting from the injection vial. For ketamine, 5-MeO-DMT succinate, 6-fluoro-DET, MDMA, and acute fluoxetine, the working solutions were prepared fresh on the day of experiment. For psilocybin, a stock solution was made and then the working solution was made from stock solution, with both solutions prepared within 1 month from the day of experiment. For chronic fluoxetine, the working solution was prepared on the first day of administration and then kept in 4°C and used for the remainder of the chronic treatment.

### Tissue collection and imaging

All the samples underwent the same tissue collection and imaging protocols. Two hours following the single-dose injection or injection of the last dose for chronic fluoxetine, mice were deeply anesthetized with isoflurane and transcardially perfused with phosphate buffered saline (P4417, Sigma-Aldrich) followed by paraformaldehyde (PFA, 4% in PBS). Brains were fixed in 4% PFA for 24 hours at 4°C, after which they were transferred to 0.1% sodium azide in PBS for storage until clearing. The SHIELD protocol was used to process the whole mouse brains. A stochastic electrotransport device^101^ was used to clear samples for 4 days at 42°C, followed by active immunolabeling using eFLASH technology integrating electrotransport^101^ and SWITCH^102^. Each brain sample was stained with 3.5 μg of rabbit anti-c-Fos monoclonal antibody (Abcam, #ab214672), followed by 10 μg of mouse anti-NeuN monoclonal antibody (Encor Biotechnology, #MCA-1B7) and then by fluorescently conjugated secondaries in 1:2 primary:secondary molar ratios (Jackson ImmunoResearch). Following active labeling, refractive index matching (n = 1.52) was done through incubation in EasyIndex (LifeCanvas Technologies). Samples were then imaged at 3.6× magnification with a SmartSPIM light sheet fluorescence microscope (LifeCanvas Technologies) at a resolution of 1.8 µm/pixel for XY sampling with 4 µm step size for Z sampling over the entire brain. Imaging was done blinded to treatment conditions.

### Atlas registration and cell counting

Fluorescence images were tile-corrected, de-striped, and registered to the Allen Brain Atlas using an automated process. For each brain, the image from the NeuN channel was registered to 8-20 atlas-aligned reference samples using SimpleElastix^103^, which implemented successive rigid, affine, and b-spline warping algorithms. The final atlas alignment value for each sample was determined by taking the average alignment generated across intermediate reference samples. Cell detection was automated by using a custom convolutional neural networked designed using the TensorFlow python package. First, a U-Net-based detection network was used to locate fluorescent puncta corresponding to c-Fos-immunolabeled cells. Second, a ResNet-based network was used to filter putative cells to arrive at a final list of cell locations. Each cell location was projected onto the Allen Brain Atlas to identify its anatomical region. We segmented the brain into 316 summary structures based on the Allen Mouse Brain Common Coordinate Framework^75^. We omitted the ‘fiber tracts’ summary structure in the analysis to focus on grey matter structures. Counts were then generated on a per-region basis for each sample.

### Batch effect correction

We observed differences in the total number of c-Fos+ cells in psilocybin samples across batch 1 and 2, saline samples across batch 1 and 2, and ketamine samples across batch 1 and 3. Batch effects are common and, in this study, may arise from differences in antibody quality, microscope condition, and/or subtle changes in the automated cell counting procedure. To correct for these differences, a scaling factor was calculated for the psilocybin, ketamine, and saline conditions individually. This factor was calculated by taking the mean total c-Fos+ cell counts of the batch 2 (psilocybin, saline) or 3 (ketamine) mice belonging to the same drug condition and dividing by mean total c-Fos+ cell counts of the batch 1 (psilocybin, saline, ketamine) mice belonging to the same drug condition. The factor was 2.78 for psilocybin, 4.94 for ketamine, and 3.11 for saline. These factors were applied to the per-region c-Fos+ cell count data in batch 1 to shift the c-Fos+ cell counts to be more comparable to the later batches. All analyses were performed after the batch effect correction. We emphasize that this batch correction step should not affect the machine learning analysis pipeline described below. This is because the first step of the pipeline is to divide per-region count by total count in each brain, meaning that the absolute values of the cell count should have minimal influence on model fits but instead it is the relative values of the cell count (e.g., proportion of c-Fos+ cell residing in one brain region over another brain region in a sample) that mattered for analysis and prediction.

### Head-twitch response

Head movements were recorded using a magnetic ear tag system as described in detail previously^33^. Briefly, an ear tag consisted of a neodymium magnet (N45, 3 mm diameter, 0.5 mm thick, #D1005-10, SuperMagnetMan) that was adhered to an aluminum ear tag (La Pias #56780, Stoelting) with cyanoacrylate glue (Super Glue Ultra Gel Control, $1739050, Loctite). The neodymium magnet was coated with a nitrocellulose marker (#7056, ColorTone) and dried for >2 h, which helped to reduce ear irritation for the mice. This magnetic ear tag was applied to the mouse’s ear using an ear tag applicator (#56791, Stoelting). For measurement, the animal was put inside a plastic cube (4” x 4” x 4”). A spool of enameled cooper wire (30 AWG) was used to wind around the cube like a solenoid, with the ends of the wire connected to a current-to-voltage preamplifier (PP444, Pyle) where the voltage was captured with a computer via a data acquisition device (USB-6001, National Instruments). Each mouse was recorded using one cube. Up to four cubes could be used to record from four mice at once inside a soundproof chamber. Data acquisition and analysis were done using custom software written in MATLAB (Mathworks). The voltage signal was sent through a 70 – 110 Hz bandpass filter because head twitch response had a characteristic ∼90 Hz frequency. The filtered signal was then processed for peak detection to identify individual head-twitch events. A protocol including parts list for the setup and the MATLAB code is available at https:// github.com/Kwan-Lab/HTR.

### Machine learning pipeline – preprocessing

The analysis pipeline used the Python package sci-kit learn (Version 1.2.1)^104^. The first step of the pipeline was preprocessing, which entails three steps: normalization, transformation, and scaling. For normalization, we divided each region’s c-Fos+ cell count by the total c-Fos+ cell count across all summary structures used. This was done to mitigate influence of batch effects across samples. For transformation, each brain region’s normalized c-Fos+ cell counts across different drug conditions were transformed using Yeo-Johnson power transformation^105^. The Yeo-Johnson transformation is a generalized form of the Box-Cox transformation. The transformation leads to data values that more closely approximate a Gaussian distribution. The Yeo-Johnson transformation was implemented in scikit-learn: *PowerTransformer(method=’yeo-johnson’, standardize=False)*. The Yeo-Johnson transformation is parameterized by one variable, lambda. The optimal lambda parameter was calculated for each brain region independently using maximum likelihood estimation to optimize for normality. For scaling, for each brain region, the *RobustScaler* module in scikit-learn was used to subtract the median value and scales values by the range of the 25^th^ to 75^th^ percentile (quartile scaling). We decided to do this, rather than subtracting mean value and standard-deviation scaling, because it is less sensitive to outliers. The c-Fos+ cell counts of each brain region after undergoing the normalization, transformation, and scaling steps are referred to as the c-Fos scores. To visualize the data, we performed dimensionality reduction on c-Fos scores across all samples using scikit-learn’s *LinearDiscriminantAnalysis* function and plotted the top two linear discriminants (**Fig. 3b**).

### Machine learning pipeline – region selection

Based on Allen Institute definition of summary structures, the brain was divided into 315 regions (316 summary structure and then ‘fiber tracts’ removed). We were concerned that a model involving c-Fos scores from 315 regions may be overfitting due to our limited sample size of 64 brains. Many regions are likely not informative and only contribute noise to the machine learning models. Therefore, we implemented a method to filter out features (i.e., the brain regions) which were not informative for distinguishing the desired drug conditions. Region selection was carried out using the Boruta algorithm, as implemented in the BorutaPy package^106^. The Boruta algorithm is an ‘all relevant features’ selection method which seeks to identify all the features with information relevant to a task. This was done by creating scrambled versions of each feature, which are called shadow features, and appending them to the original data set. This expanded data set was then used to fit a random forest classifier, as implemented in scikit-learn. We used the BorutaPy package to automatically select the number of trees for the *RandomForestClassifier()* module based on the size of the feature set. Following this, a threshold was established based on the highest feature importance amongst shadow features. Features exceeding this threshold were considered ‘hits’ and recorded. This procedure was repeated 100 times. The distribution across these 100 iterations created a binomial distribution. The BorutaPy package rejected features based on the cumulative distribution function of a binomial distribution where p = 0.5, alpha = 0.05, and n = number of hits. Features (i.e., brain regions) that were not rejected by this criterion were the feature included for the next stage of the pipeline.

### Machine learning pipeline – classification

We used the c-Fos scores of the selected brain regions to fit a ridge regression model (L2 normalized logistic regression). The regularization parameter C is a hyperparameter used to modulate the penalty strength. Given the interconnected nature of the exact feature set and hyperparameter, as well as our desire to eventually merge results across many cross-validation splits of the data, we opted to fix this parameter to its default value of 1. The ‘multinomial’ setting was used to generalize from binary classification to multi-class classification.

### Cross validation to determine prediction accuracy

The data were evaluated using the aforementioned pipeline using 4-fold splits, where 75% of the data (i.e., 6 brain samples) in each drug condition was used to train and fit the model, while the remaining 25% of the data (i.e., 2 brain samples) was used to test the model. Importantly, preprocessing parameters (e.g., lambda in Yeo-Johnson transformation) and feature selection (brain regions to be included) were chosen using only the training data to ensure no data leakage. Nevertheless, after those stages were fixed, the test data would undergo the same preprocessing and feature selection steps before being inputted into the ridge regression model to generate the prediction of the drug condition. We performed 100 iterations, each time using a randomized splits for each drug condition, generated by scikit-learn’s *StratifiedShuffleSplit()* function. Combining the outcome across the 100 iterations, the predicted classifications were used to generate a mean confusion matrix (**Fig. 3c**). The probabilities assigned to each label for each test data point were combined to create a composite precision recall curve, generated using scikit-learn’s *precision_recall_curve* function (**Fig. 3d**, **Fig. 4b**). The scikit-learn’s *auc* function was used to calculate the area under the curve for each composite precision recall curve (legend of **Fig. 3d**). We used numpy’s random seeds and state objects (numpy.random.RandomState()) to generate reproducible results. The cross validation splitting function was seeded with an integer, per scikit-learn’s recommendations. Remaining random states were set using a random state object. A null distribution for area under the precision recall curve was established by shuffling labels during each cross validation split prior to model fitting and label prediction (**Fig. 4b**).

### Shapley additive explanation

SHAP values were generated by the *LinearExplainer* object from the SHAP package, which accepted test data points and the fit model. We set the feature perturbation parameter of the *LinearExplainer* to ‘*correlation_dependent*’. SHAP values were generated in part by breaking dependencies across features and testing the influence of perturbations on individual features. This ran the risk of creating unrealistic feature combinations, because many brain regions which would normally change in lockstep may be changed individually by the algorithm to infer feature importance, which would lead to inflated feature importance scores^107^. By using the “correlation dependent” intervention, additional measures were taken to address correlations in the feature space and credit was distributed more appropriately. The SHAP values for each test data point were combined across the data splits from the 100 iterations to arrive at composite SHAP summary plots (**Figs. 5c, 6a, 6c, 6e**). We determined which brain regions were included in >=75% of the cross-validation splits of the data (**Figs. 4c, 4d**). Regions meeting this criterion were visualized using the brainrender package^108^ (**Figs. 5d, 6b, 6d, 6f**).

### Leave-one-drug-out analysis

The fitting of the pipeline (*pipelineObj.fit*) was performed on a reduced dataset of cFos scores, excluding all samples in the 6-fluoro-DET condition. That is, for each split, training data were c-Fos+ cell count from 75% of the samples from 7 conditions (psilocybin, ketamine, 5-MeO-DMT, MDMA, acute fluoxetine, chronic fluoxetine, and saline). The test data consist of c-Fos+ cell count from the remaining 25% of the samples from those 7 conditions and 25% of the samples drawn from the left-out condition of 6-fluoro-DET. For linear discriminant analysis, the full dataset was transformed (*pipelineObj.transform*) and plotted using multiple calls to the seaborn scatterplot function (*sns.scatterplot*).

