## Supplementary Figures for "Classification of psychedelics and psychoactive drugs based on brain-wide imaging of cellular c-Fos expression"

**Supplementary Figure 1-2**

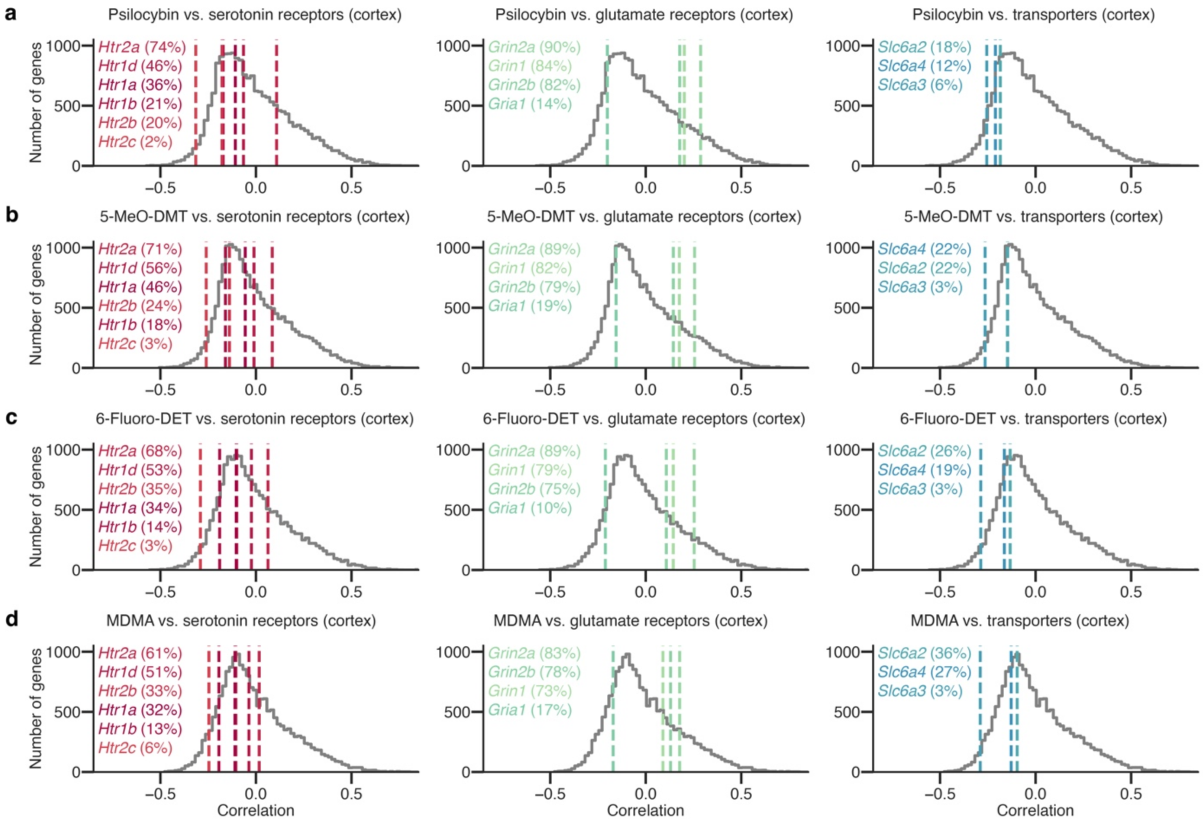


**Supplementary Figure 1**: **Exploratory analysis of the potential receptors contributing to drug-evoked c-Fos expression. a.** Correlation coefficients computed for psilocybin condition using regions across the entire brain. Colored lines, correlation coefficients for select serotonin receptor genes (left), glutamate receptor genes (middle), and monoamine transporter genes (right). The percentile values are indicated in parentheses. Gray line, histogram of correlation coefficients for 19,413 genes in the mouse genome. **b.** Similar to a for 5-MeO-DMT. **c.** Similar to a for 6-F-DET. **d.** Similar to a for MDMA.

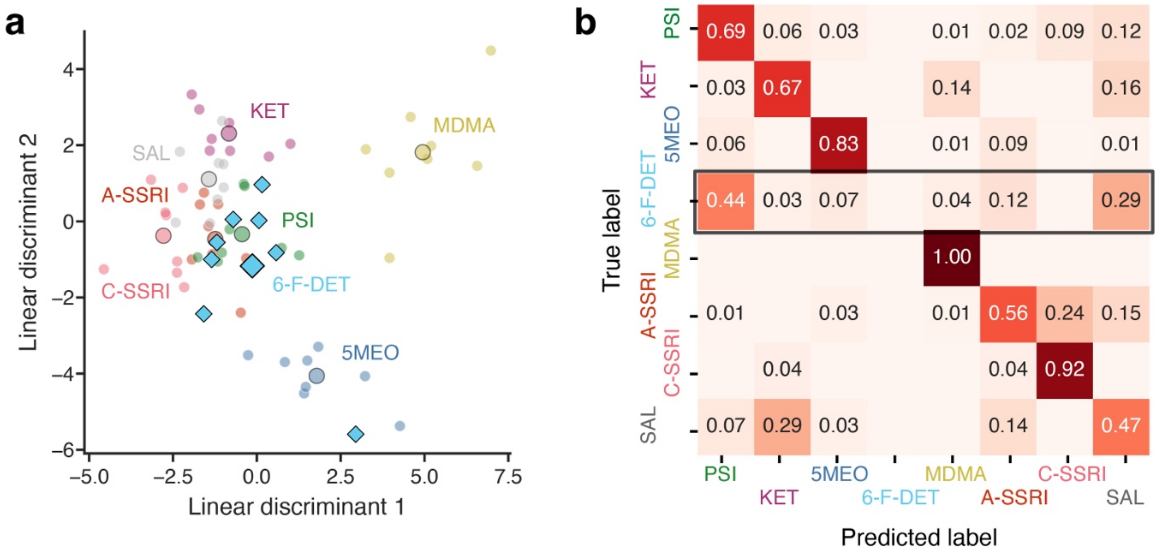


**Supplementary Figure 2: A machine learning pipeline for drug prediction and performance of one-versus-rest classification, trained using 7 conditions for a leave-one-drug-out analysis.** The pipeline is the same as Fig. 3a, except that the training data are c-Fos+ cell count from 75% of the samples from 7 conditions (psilocybin, ketamine, 5-MeO-DMT, MDMA, acute fluoxetine, chronic fluoxetine, and saline). The test data consist of c-Fos+ cell count from the remaining 25% of the samples from those 7 conditions and 25% of the samples drawn from the left-out condition of 6-fluoro-DET. **a,** Linear discriminant analysis of the c-Fos scores to visualize the data in a low dimensional space. **b,** The confusion matrix showing the mean proportion of predicted labels for each of the true labels across all splits. Note that column for 6-F-DET as a predicted label is blank because the classifier was not trained on the 6-F-DET data and so would not predict that label.
